# A Unified Spatial AI Framework for Cross-Domain Tissue-State Analysis in Trauma, Oral, and Cardiovascular Pathology

**DOI:** 10.64898/2026.06.07.730662

**Authors:** Tuan D. Pham

**Affiliations:** Barts and The London School of Medicine and Dentistry, Queen Mary University of London, Turner Street, E1 2AD, London, UK

**Keywords:** Artificial intelligence, data science, spatial transcriptomics, trauma–oral–cardiovascular axis, cross-domain inflammatory biology

## Abstract

**Objective:** To develop a cross-domain spatial AI framework for identifying conserved tissue-state organisation across trauma, oral disease, and cardiovascular tissue using spatial transcriptomic data.

**Methods:** Four public spatial transcriptomic datasets spanning wound healing, periodontitis, oral squamous cell carcinoma, and cardiac tissue were integrated using recurrence modelling, graph-based spatial learning, fuzzy tissue-state analysis, and tensor decomposition. Cross-domain coupling, spatial fragmentation, recurrence structure, and permutation-based topological validation were evaluated.

**Results:** Six conserved fuzzy tissue states were identified, dominated by extracellular matrix remodelling, fibroblast/stromal activation, endothelial signalling, and inflammatory pathways. Latent embedding analysis demonstrated strong overlap between trauma and oral domains, while cardiovascular tissue exhibited more compact spatial organisation. Oral inflammatory tissue showed the highest fragmentation, whereas cardiovascular tissue demonstrated greater recurrence coherence. Tensor decomposition identified conserved stromal-remodelling programmes across domains. Permutation testing confirmed significantly elevated graph modularity and reduced spatial entropy relative to null distributions.

**Conclusion:** The proposed framework identified conserved spatial tissue-state architecture linking wound healing, oral pathology, and cardiovascular tissue despite differences in tissue origin, pathology, and acquisition technology.

**Significance:** These findings demonstrate the potential of spatial AI for investigating conserved stromal and inflammatory microenvironmental organisation across clinically related disease systems and may support spatial biology research in trauma–oral–systemic health.

## 1 Introduction

Trauma care, oral inflammatory disease, oral cancer, and cardiovascular pathology are major contributors to global morbidity and healthcare burden [1, 2]. Although these conditions affect anatomically distinct tissues, increasing evidence suggests that they share common inflammatory, stromal, vascular, and extracellular matrix (ECM) remodelling mechanisms that regulate tissue injury, repair, fibrosis, and chronic disease progression [3, 4, 5]. Chronic oral inflammatory conditions such as periodontitis have been associated with systemic cardiovascular dysfunction, endothelial injury, and persistent inflammatory activation [2, 6, 7], while post-traumatic tissue repair and tumour-associated stromal remodelling exhibit overlapping fibroblast activation and angiogenic processes [8, 9]. Understanding how these biological programmes are spatially organised across tissues is therefore important for improving mechanistic understanding of trauma-oral-systemic disease relationships, tissue repair biology, and translational precision medicine [2, 10].

Recent advances in spatial transcriptomics have enabled high-resolution investigation of tissue microenvironments and spatial cellular organisation [10, 11]. Technologies such as 10x Genomics Visium permit simultaneous analysis of gene expression and tissue architecture, facilitating identification of spatially localised biological programmes within intact tissue sections [10, 12]. Concurrently, graph neural networks, manifold learning, recurrence-based representations, tensor decomposition methods, and multi-modal data integration have increasingly been applied to characterise tumour ecosystems, inflammatory niches, wound healing, and cardiovascular tissue remodelling [13, 14, 15, 16, 17, 18]. These approaches have demonstrated the utility of graph-based modelling for spatial cellular interactions [14, 15], recurrence analysis for capturing nonlinear biological organisation [19, 20], and tensor methods for identifying latent multi-domain biological programmes [16, 21]. More broadly, computational biomedical engineering has increasingly adopted topology-aware, graph-theoretic, and higher-order representation learning frameworks for analysing complex and heterogeneous biomedical data [22, 23, 24]. Prior studies have shown that multi-graph learning, network-constrained modelling, and latent tensor representations can reveal structured biological patterns across heterogeneous datasets [22, 23]. However, most existing spatial artificial intelligence (AI) studies remain focused on single diseases, isolated tissue systems, or supervised classification tasks, limiting their ability to identify conserved spatial biological organisation across multiple pathological domains [15, 14].

Despite these advances, several important limitations remain. Current approaches rarely investigate whether conserved spatial tissue-state organisation exists across biologically distinct but clinically related disease systems such as trauma-related tissue repair, oral disease, and cardiovascular pathology [17, 18, 8, 25]. Many existing methods rely on rigid clustering or discrete cell-state assignment, conventional classification and image segmentation methods, which may inadequately represent the continuous and overlapping biological transitions commonly observed in inflammatory and stromal tissue microenvironments [15, 26]. Furthermore, spatial AI frameworks frequently emphasise predictive classification performance rather than unsupervised discovery of conserved biological organisation across heterogeneous datasets [14, 15]. Existing studies also often lack rigorous topological validation against permutation-derived null models, limiting interpretation of whether observed spatial structures reflect meaningful biological organisation or random clustering artefacts. Finally, few studies integrate recurrence modelling, graph-constrained learning, and tensor decomposition within a unified framework for analysing cross-domain spatial transcriptomic organisation [16, 21, 15].

To address these limitations, this study presents a unified cross-domain spatial AI framework integrating recurrence modelling, graph-based spatial learning, fuzzy tissue-state analysis, and tensor decomposition to identify conserved spatial biological programmes across trauma, oral, and cardiovascular tissues. The framework models continuous and overlapping tissue states using fuzzy latent representations and incorporates cross-domain coupling, spatial fragmentation, recurrence analysis, and permutation-based validation to quantify conserved and non-random spatial organisation.

The study makes several contributions. First, it introduces a cross-domain spatial AI framework integrating recurrence modelling, graph-based spatial learning, fuzzy tissue-state analysis, and tensor decomposition for analysing heterogeneous spatial transcriptomic datasets. Second, the framework identifies conserved latent tissue-state organisation linking trauma, oral inflammatory disease, oral pathology, and cardiovascular tissue. Third, the results demonstrate that ECM remodelling, fibroblast activation, endothelial signalling, and inflammatory pathways constitute dominant conserved spatial programmes across biological domains. Fourth, the framework introduces fuzzy tissue-state representations capable of modelling continuous and overlapping biological transitions within stromal-inflammatory microenvironments. Finally, the study incorporates quantitative analysis of cross-domain coupling, spatial fragmentation, local recurrence organisation, latent pathway–state–domain programmes, and permutation-based topological validation to evaluate whether the identified spatial organisation exceeds random tissue-state structure.

## 2 Methods

### 2.1 Cross-domain study design and public spatial transcriptomic datasets

This study was designed as a cross-domain spatial biology framework to investigate whether trauma, oral disease, and cardiovascular pathology share conserved spatial tissue states and latent biological programmes. The central hypothesis is that these clinically related conditions exhibit recurrent inflammatory, stromal, vascular, hypoxic, ECM, immune, and metabolic tissue programmes that can be identified across independent public spatial transcriptomic datasets. The study does not assume direct causality between disease domains. Instead, it evaluates whether public spatial omics data provide reproducible evidence of conserved tissue-state organisation across biologically connected pathological conditions.

Because the datasets originate from different tissues, studies, and experimental platforms, downstream analyses focused on latent pathway-level and tissue-state representations rather than direct gene-level comparisons. This strategy reduces dataset-specific technical variation and tissue-related bias while enabling cross-domain modelling of conserved biological programmes.

The trauma domain was represented by the human wound-healing spatial transcriptomic dataset GSE241124 [27]. This dataset contains spatial transcriptomic profiles from human skin wound samples collected across multiple temporal stages of wound healing, including inflammatory, proliferative, and remodelling phases. Although the tissue source is skin, the dataset provides a biologically relevant model of acute tissue injury, inflammation, stromal activation, angiogenesis, and ECM remodelling.

The oral-disease domain was represented using two independent datasets. The first dataset, GSE206621 [28], contains human gingival and oral mucosal tissue sections from healthy and periodontitis-affected individuals analysed using 10x Genomics Visium spatial transcriptomics. This dataset was used to model chronic oral inflammatory tissue organisation and immune–stromal interactions within the oral microenvironment. The second dataset, GSE208253 [29], contains spatial transcriptomic profiles from oral squamous cell carcinoma (OSCC) tissue regions, including tumour-core and invasive-margin regions. This dataset was included because oral cancer tissue exhibits inflammatory, stromal, epithelial, vascular, and hypoxic remodelling programmes relevant to oral-systemic disease biology and pathological tissue reorganisation.

The cardiovascular domain was represented by the human cardiac spatial transcriptomic dataset GSE135805 [30], which was generated alongside a plasma proteomic characterisation of human heart failure, and contains spatial transcriptomic profiles from healthy and diseased human cardiac tissue, including cardiomyopathic remodelling states. This dataset was used to model inflammatory, vascular, stromal, fibrotic, and metabolic tissue states associated with cardiovascular pathology and cardiac remodelling.

The biological domains were therefore defined as *D* = {*T, O, C*}, where *T, O*, and *C* denote trauma, oral disease, and cardiovascular pathology, respectively.

### 2.2 Gene harmonisation and normalisation

Let 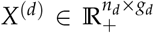 denote the spatial transcriptomic expression matrix for dataset *d, d* = 1, · · ·, *D*, where *n*_*d*_ is the number of spatial locations and *g*_*d*_ is the number of measured genes. The corresponding spatial coordinate matrix is denoted by 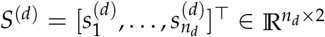, where 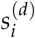 represents the spatial coordinate of location *i*. To enable cross-dataset integration, a common gene set was defined as

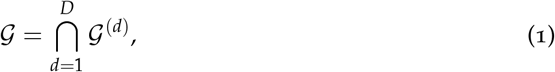

where 𝒢 ^(*d*)^ denotes the set of genes present in dataset *d*. Gene identifiers were harmonised across datasets by mapping Ensembl gene identifiers (stable gene IDs assigned by the Ensembl genome annotation system [31]) to HGNC (HUGO Gene Nomenclature Committee [32]) gene symbols using a curated mapping table. Only genes shared across all datasets were retained for downstream analyses.

For each spatial location *i* and gene *j*, library-size normalisation was applied using

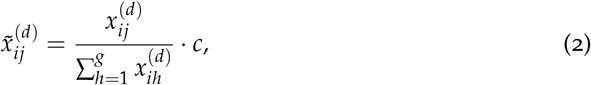

where 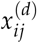 denotes the raw expression value and *c* is a scaling constant fixed to *c* = 10^4^.

The normalised expression matrix is denoted by

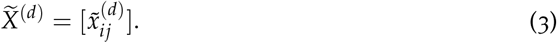

### 2.3 Spatial graph construction

Each dataset was represented as a weighted graph:

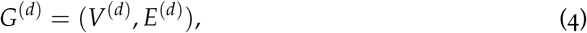

where *V*^(*d*)^ denotes the set of spatial locations and *E*^(*d*)^ denotes the set of spatial neighbourhood edges.

Spatial neighbourhoods were constructed using a *k*-nearest neighbour graph with *k* = 8 to capture local spatial organisation while avoiding overly dense graphs. The edge weight between locations *i* and *j* was defined using a Gaussian similarity kernel:

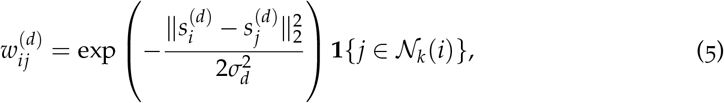

where *N*_*k*_(*i*) denotes the set of *k*-nearest neighbours of location *i, σ*_*d*_ is the Gaussian kernel bandwidth parameter, and **1**{·} denotes the indicator function. The bandwidth parameter was estimated as the median neighbour distance within each dataset, providing a data-adaptive scale robust to different spatial resolutions.

The weighted adjacency matrix is denoted by

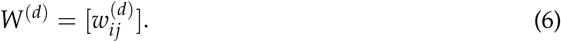

The diagonal degree matrix is

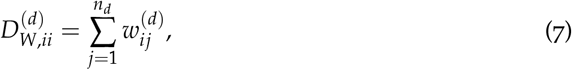

and the graph Laplacian is

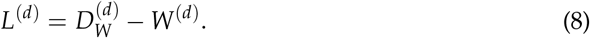

The graph Laplacian regularised the spatial embedding process by encouraging neighbouring tissue regions with strong graph connectivity to exhibit similar latent representations.

### 2.4 Pathway activity scoring

To improve biological interpretability, gene expression was projected onto pathway-level activity scores. Let 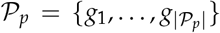 denote the gene set associated with pathway *p*, where *p* = 1, …, *P*. Only pathways containing at least *g*_min_ = 3 genes remaining after cross-dataset harmonisation were retained for downstream analyses. Curated biological pathways were obtained from Gene Matrix Transposed (GMT) pathway collections, a standard file format for representing predefined gene sets and biological pathways [33]. Each GMT entry contains a pathway identifier followed by the associated member genes.

For spatial location *i*, pathway activity was estimated as

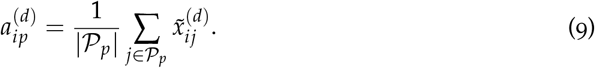

The resulting pathway activity matrix is 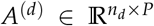, where rows and columns correspond to spatial locations and pathway activities, respectively.

### 2.5 Spatial latent representation learning

Low-dimensional spatial latent representations were learned from the normalised expression matrix using iterative graph-based spatial smoothing. Let 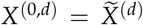 denote the initial normalised expression matrix for dataset *d*. Spatial smoothing was iteratively performed as

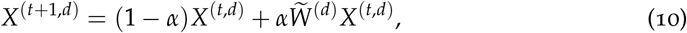

Where

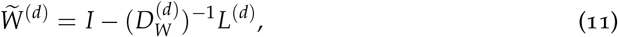

is the Laplacian-derived diffusion operator, which enforces neighbouring spatial locations to exhibit similar representations while preserving local tissue topology. The smoothing weight was fixed to *α* = 0.15 and applied for 10 iterations.

After smoothing, principal component analysis was applied to obtain the latent embedding matrix

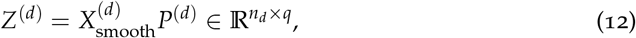

where 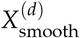 denotes the final smoothed expression matrix and *P*^(*d*)^ contains the first *q* principal component loading vectors. The latent dimension was fixed to *q* = 20 to retain the principal expression variation while reducing noise and computational complexity.

### 2.6 Fuzzy tissue-state modelling

The learned embeddings from all datasets were pooled as

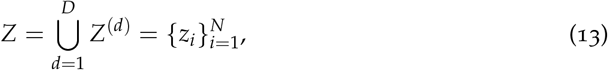

Where

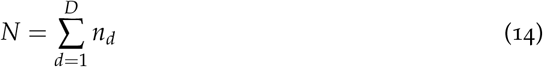

is the total number of spatial locations.

The fuzzy *c*-means clustering (FCM) algorithm [34] was applied to identify overlapping latent tissue states. Let

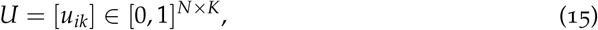

denote the fuzzy membership matrix, where *u*_*ik*_ represents the partial membership of spatial location *i* in tissue state *k*, with *k* = 1, …, *K*. The number of tissue states was fixed to *K* = 6 to capture overlapping stromal, inflammatory, vascular, and transitional microenvironmental programmes while preserving smooth latent tissue-state organisation. The memberships satisfy

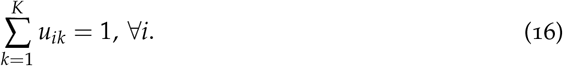

The FCM objective function is defined as

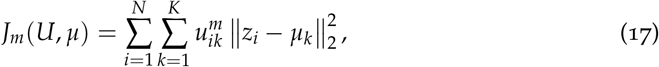

where *μ*_*k*_ ∈ ℝ^*q*^ denotes the centroid of tissue state *k*, and *m >* 1 is the fuzzifier exponent. The fuzzifier was fixed to the commonly used value *m* = 2. After initialising *U*, the cluster centres *μ*_*k*_ and membership values *u*_*ik*_ were iteratively updated until convergence or a maximum of 100 iterations was reached.

Tissue-state labels were subsequently assigned as

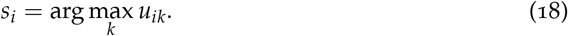

Exponentiated memberships were additionally computed as

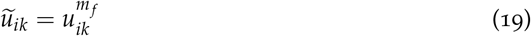

for downstream pathway and conservation analyses. Exponentiated memberships were used to emphasise high-confidence tissue-state assignments while reducing the influence of ambiguous boundary regions.

### 2.7 Cross-domain tissue-state conservation

For tissue state *k*, the contribution of dataset *d* was computed as

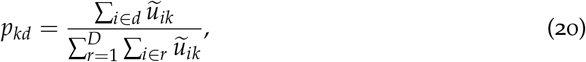

where *p*_*kd*_ represents the proportion of tissue state *k* contributed by dataset *d*.

The entropy-based conservation score was defined as

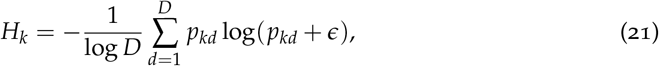

where *ϵ* is a small value used to avoid log(0). Values of *H*_*k*_ close to one indicate tissue states broadly conserved across multiple datasets and domains, whereas lower values indicate stronger domain specificity.

### 2.8 Trauma–oral–cardiovascular coupling

The three biological domains are denoted by *D*_*T*_, *D*_*O*_, and *D*_*C*_, corresponding to trauma, oral disease, and cardiovascular pathology, respectively.

For each domain *r*, the fuzzy tissue-state distribution was computed as

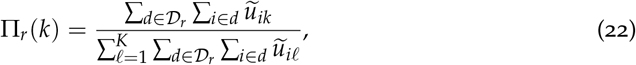

yielding the tissue-state composition vector

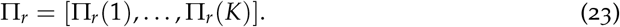

The similarity between domains *r* and *s* was measured using the Jensen–Shannon similarity [35]:

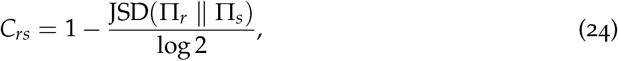

which denotes the off-diagonal coupling between domains *r* and *s*, and where JSD(·) denotes the Jensen–Shannon divergence.

The global coupling score was defined as

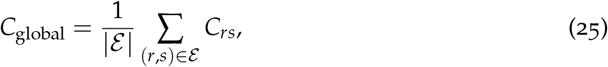

where

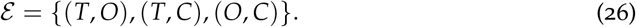

### 2.9 Spatial recurrence and tissue fragmentation

Spatial recurrence analysis was performed within latent embedding space. A continuous-valued spatial recurrence score between neighbouring spatial locations *i* and *j* was defined as

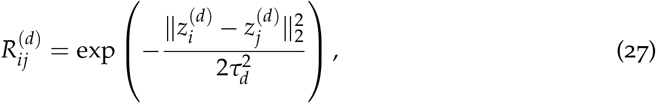

where *τ*_*d*_ is a dataset-specific recurrence scale parameter estimated from median neighbourhood distances within latent space.

The local recurrence density at spatial location *i* was computed as

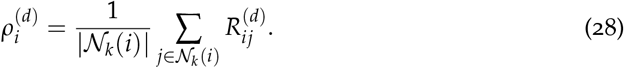

The pathological fragmentation index was defined as

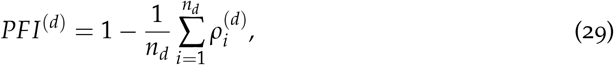

which quantifies the degree of spatial fragmentation of latent tissue organisation. Since 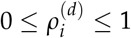, the index satisfies 0 ≤ *PFI*^(*d*)^ ≤ 1, where values close to zero indicate highly coherent spatial organisation and values close to one indicate highly fragmented tissue structure.

### 2.10 Tensor decomposition of conserved biological programmes

To identify interpretable cross-domain programmes, a pathway–state–domain tensor was constructed as

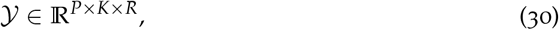

where *P* is the number of pathways, *K* is the number of tissue states, and *R* = 3 indicates the number of biological domains corresponding to trauma, oral, and cardiovascular tissue domains, such that *D* = {*T, O, C*}.

Tensor elements were defined as

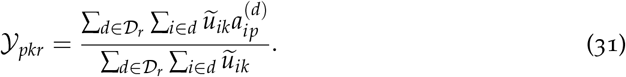

Canonical polyadic (CP) decomposition was subsequently applied:

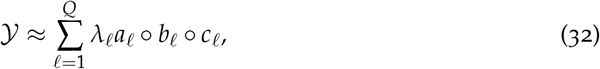

where *Q* denotes tensor rank, which was fixed to 3, *λ*_*ℓ*_ denotes the component weight, and º denotes the vector outer product.

Canonical polyadic (CP) decomposition was subsequently applied:

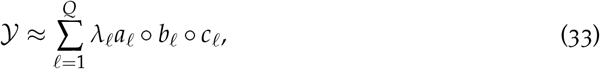

where *Q* denotes the tensor rank, fixed to *Q* = 3, *λ*_*ℓ*_ denotes the component weight, *a*_*ℓ*_, *b*_*ℓ*_, and *c*_*ℓ*_ contain pathway, tissue-state, and domain loadings, respectively, and º denotes the vector outer product. Larger loading values indicate stronger contribution of the corresponding pathway, tissue state, or domain to the latent biological programme represented by component *ℓ*. Components exhibiting elevated domain loadings across multiple domains were interpreted as conserved biological programmes linking trauma, oral disease, and cardiovascular pathology.

### 2.11 Topological validation against permuted controls

Topological validation was performed by comparing observed tissue-state organisation against spatially permuted controls. Graph modularity was computed as

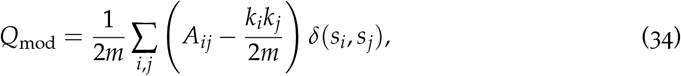

where *A*_*ij*_ denotes the (*i, j*)-th entry of the weighted graph adjacency matrix constructed from the *k*-nearest neighbour spatial graph, *δ*(·, ·) is the Kronecker delta function, and

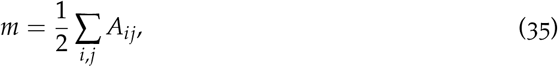

denotes the total graph edge weight. The weighted node degree is defined as

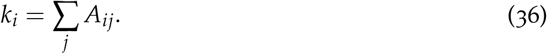

Neighbourhood entropy was computed as

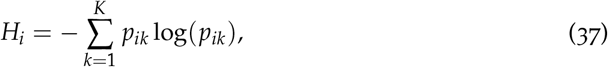

where *p*_*ik*_ denotes the proportion of neighbouring nodes belonging to tissue state *k*. Dataset-level spatial entropy was then estimated as

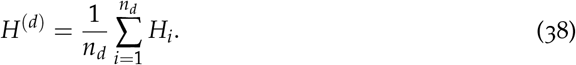

Permutation validation independently randomised (i) tissue-state labels for topological analyses and (ii) latent embeddings for recurrence-based analyses while preserving graph topology. Latent embedding permutation was defined as

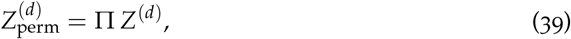

where Π denotes a random permutation operator. Permutation testing used 1000 random permutations to provide a practical empirical null distribution while maintairather than exhaustive pairwise node comparisons, enabling scalable validation on large spatial graphs.

## 3 Results

### 3.1 Integrated latent tissue-state organisation across biological domains

The integrated spatial biology framework analysed four datasets spanning trauma, oral inflammation, oral pathology, and cardiovascular tissue (Table 1). Following preprocessing, pathway harmonisation, graph construction, and latent embedding, a total of 106,377 spatial observations were retained for cross-domain analysis.

**Table 1:** Summary of datasets included in the study.

| Dataset ID | Biological domain | Tissue context | Technology | Spatial observations |
| --- | --- | --- | --- | --- |
| GSE241124 | Trauma | Human wound healing | Visium ST | 25,969 |
| GSE206621 | Oral inflammation | Periodontitis/oral mucosa | Visium ST | 34,204 |
| GSE208253 | Oral pathology | OSCC | Visium ST | 32,103 |
| GSE135805 | Cardiovascular | Human cardiac tissue | Spatial Transcriptomics | 7,504 |

The latent embedding demonstrated substantial overlap between trauma, oral, and cardiovascular domains, indicating conserved spatial biological organisation across tissue contexts (Figure 1(a)). Trauma and oral tissues occupied strongly overlapping latent regions, whereas cardiovascular tissue formed a comparatively compact central distribution within the shared manifold.

**Figure 1:**
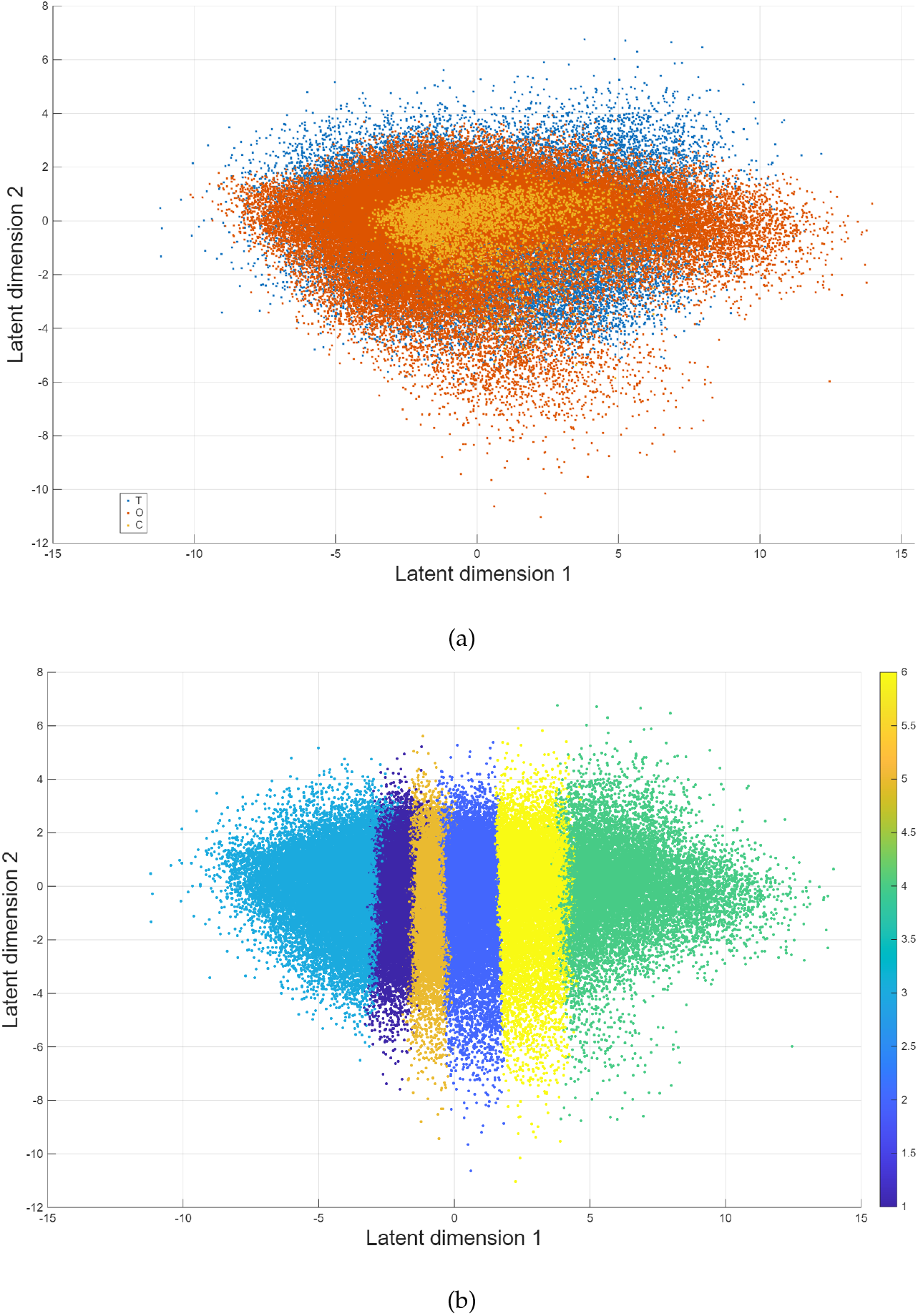
Latent tissue-state organisation across biological domains: latent embedding coloured by biological domain, showing substantial overlap between trauma (blue), oral (orange), and cardiovascular (yellow) tissues (a); and fuzzy tissue-state assignments demonstrating six overlapping latent tissue states distributed along the latent manifold (b).

Fuzzy tissue-state modelling (Figure 1(b)) identified six latent tissue states distributed continuously along the latent space rather than forming sharply separated clusters. The substantial overlap between colours reflects continuous and transitional spatial microenvironmental organisation rather than discrete tissue classes. The gradual transitions between neighbouring states further support the presence of continuous biological progression and overlapping stromal–inflammatory microenvironments.

### 3.2 Conserved tissue-state pathways and cross-domain coupling

The six fuzzy tissue states were characterised by recurrent pathway activation patterns dominated by ECM remodelling, fibroblast/stromal activation, endothelial signalling, and inflammatory activity (Table 2). States 2, 4, and 6 exhibited particularly strong stromal and ECM-associated signatures, whereas States 1 and 3 showed relatively stronger vascular and inflammatory enrichment.

**Table 2:** Dominant pathway annotations associated with fuzzy tissue states.

| State | Top pathway | Secondary pathway | Biological interpretation |
| --- | --- | --- | --- |
| 1 | ECM remodelling | Endothelial/vascular activity | Vascular remodelling state |
| 2 | ECM remodelling | Fibroblast/stromal activation | Stromal repair state |
| 3 | ECM remodelling | Inflammatory signalling | Inflammatory remodelling state |
| 4 | ECM remodelling | Fibroblast/stromal activation | Matrix-producing stromal state |
| 5 | ECM remodelling | Endothelial/vascular activity | Angiogenic remodelling state |
| 6 | ECM remodelling | Fibroblast/stromal activation | Activated fibro-inflammatory state |

State conservation entropy analysis demonstrated broad cross-domain recurrence of all tissue states (Table 3). Entropy values ranged from 0.763 to 0.900, with a mean entropy of 0.860, indicating that most tissue states were distributed across multiple biological domains rather than remaining domain-specific.

**Table 3:** State conservation entropy and relative domain contributions. T = Trauma, O = Oral, and C = Cardiovascular.

| State | Entropy | T (%) | O (%) | C (%) |
| --- | --- | --- | --- | --- |
| 1 | 0.900 | 23.4 | 71.7 | 4.9 |
| 2 | 0.889 | 24.1 | 71.6 | 4.3 |
| 3 | 0.763 | 29.6 | 66.5 | 3.9 |
| 4 | 0.841 | 26.3 | 69.7 | 4.0 |
| 5 | 0.872 | 25.4 | 70.5 | 4.1 |
| 6 | 0.894 | 24.0 | 71.8 | 4.2 |

Pairwise domain coupling further demonstrated strong similarity between tissue-state organisations across biological domains (Figure 2; Table 4). The strongest coupling was observed between trauma and oral tissues (0.994), whereas cardiovascular tissue demonstrated slightly lower but still substantial coupling with both domains.

**Table 4:** Cross-domain coupling matrix derived from latent tissue-state organisation.

|  | Trauma | Oral | Cardiovascular |
| --- | --- | --- | --- |
| Trauma | 1.000 | 0.994 | 0.945 |
| Oral | 0.994 | 1.000 | 0.940 |
| Cardiovascular | 0.945 | 0.940 | 1.000 |

**Figure 2:**
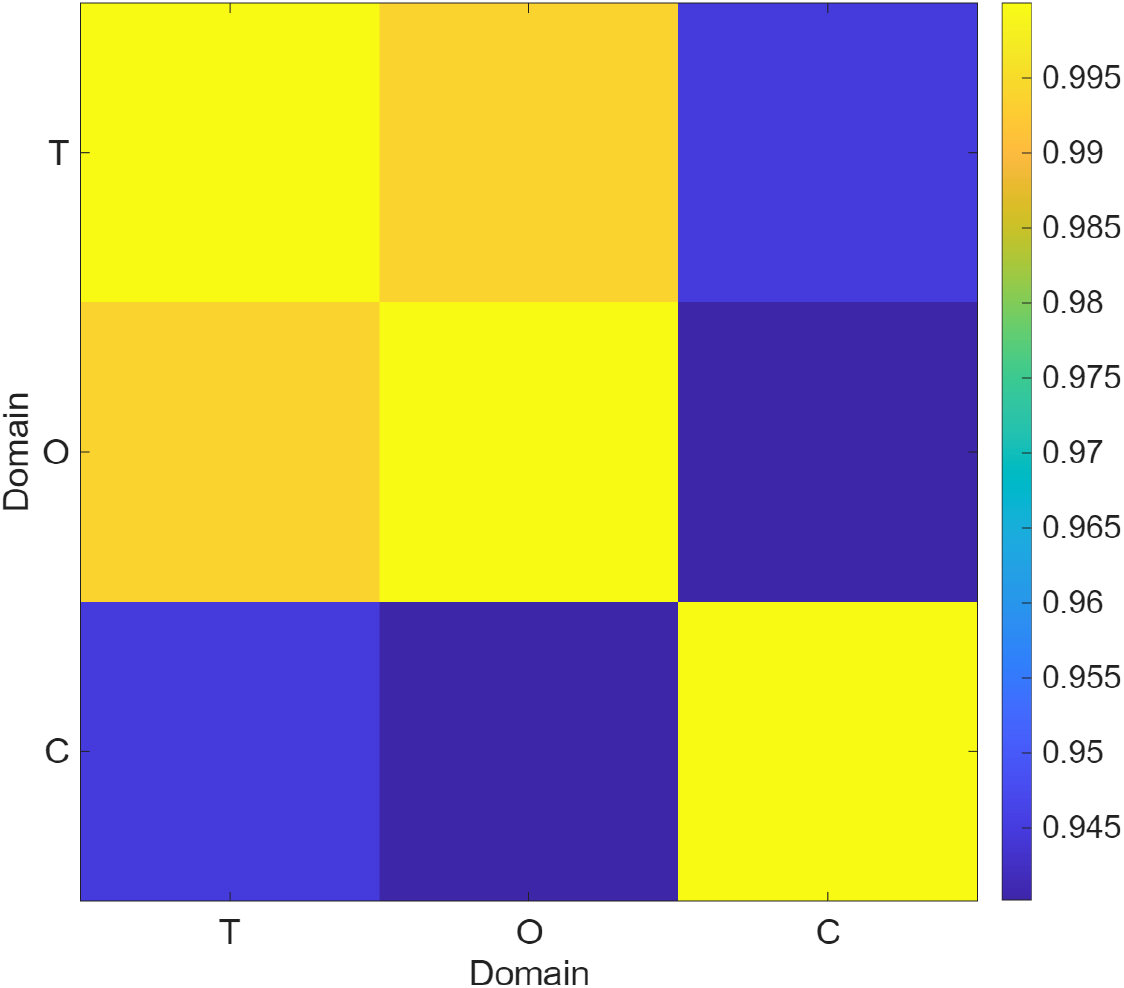
Cross-domain coupling matrix derived from latent tissue-state organisation.

It should be noted that the cardiovascular domain contributed fewer spatial observations (7,504) relative to the trauma and oral datasets (25,969—34,204), resulting in comparatively lower cardiovascular representation across all tissue states (3.9–4.9%). Although cross-domain coupling remained substantial, this imbalance should be considered when interpreting the degree of cardiovascular conservation reported here. Collectively, these findings indicated strong conservation of stromal and inflammatory tissue-remodelling programmes across trauma, oral disease, and cardiovascular tissue.

### 3.3 Spatial tissue-state organisation and recurrence structure

Spatial mapping of fuzzy tissue states revealed heterogeneous yet spatially coherent tissue-state organisation across datasets. For example, OSCC tissue (GSE208253) demonstrated regionally coherent tissue-state structure with distinct spatial domains (Figure 3(a)).

**Figure 3:**
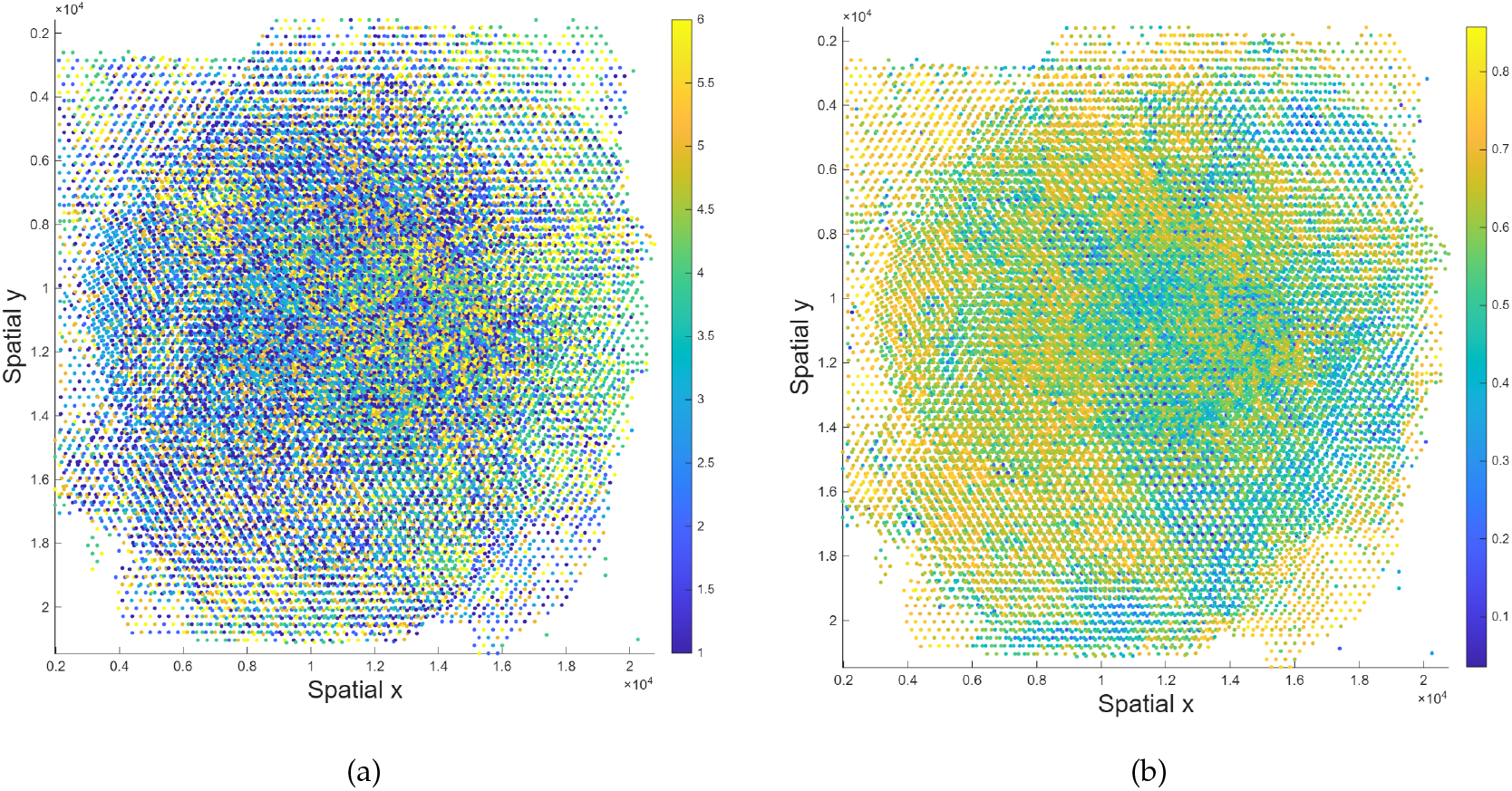
GSE208253: Spatial distribution of fuzzy tissue states (a), and local recurrence density map (b).

These spatial differences were quantitatively reflected in the fragmentation and recurrence metrics summarised in Table 5. Oral inflammatory tissue demonstrated the highest PFI, whereas cardiovascular tissue exhibited the lowest fragmentation and spatial entropy, indicating comparatively stronger spatial coherence.

**Table 5:**
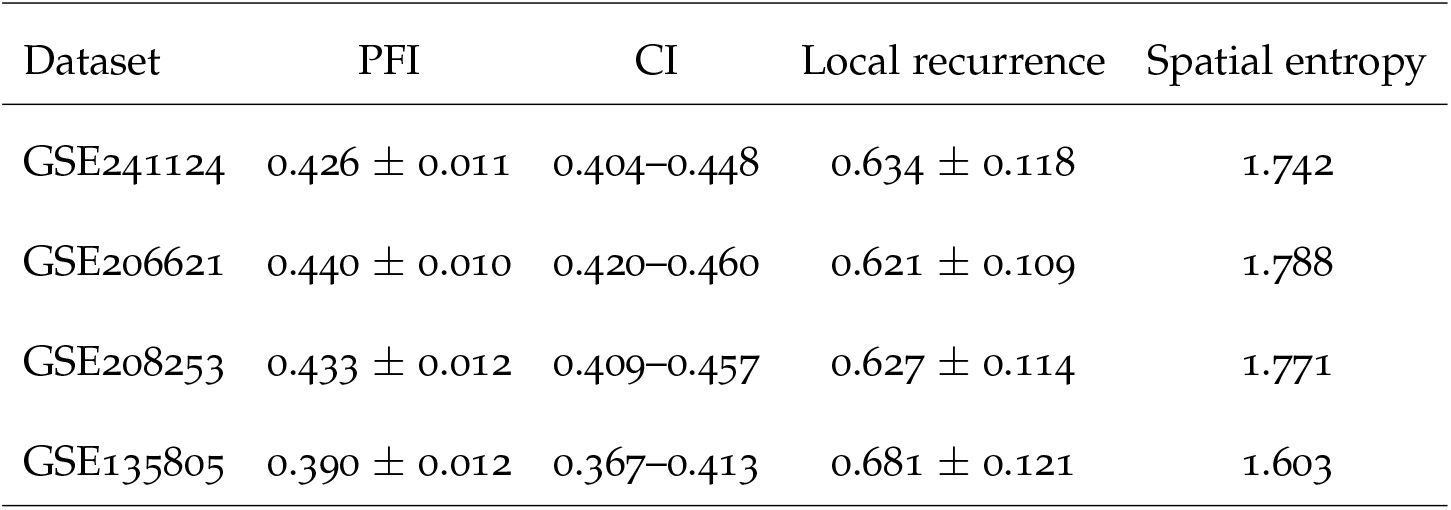
Spatial fragmentation and recurrence metrics. Values are reported as mean *±* SD where applicable. CI denotes the 95% confidence interval.

| Dataset | PFI | CI | Local recurrence | Spatial entropy |
| --- | --- | --- | --- | --- |
| GSE241124 | 0.426 ± 0.011 | 0.404–0.448 | 0.634 ± 0.118 | 1.742 |
| GSE206621 | 0.440 ± 0.010 | 0.420–0.460 | 0.621 ± 0.109 | 1.788 |
| GSE208253 | 0.433 ± 0.012 | 0.409–0.457 | 0.627 ± 0.114 | 1.771 |
| GSE135805 | 0.390 ± 0.012 | 0.367–0.413 | 0.681 ± 0.121 | 1.603 |

Local recurrence density maps further demonstrated distinct recurrence organisation across inflammatory and pathological tissues. In particular, GSE208253 exhibited relatively preserved high-density recurrence regions and stronger spatial coherence (Figure 3(b)).

### 3.4 Tensor-derived conserved programmes and topological validation

CP tensor decomposition identified three dominant pathway–state–domain programmes shared across biological domains (Table 6). The dominant component was characterised by ECM remodelling and fibroblast activation distributed across trauma, oral, and cardiovascular tissues, whereas secondary components reflected vascular and inflammatory programmes.

**Table 6:** Tensor-derived cross-domain biological programmes.

| Component | Dominant pathways | Dominant states | Domain loading pattern |
| --- | --- | --- | --- |
| 1 | ECM remodelling, fibroblast activation | 2, 4, 6 | Strong T-O-C coupling |
| 2 | Endothelial/vascular activity, angiogenesis | 1, 5 | Moderate T-O-C coupling |
| 3 | Inflammatory signalling, stress response | 3, 6 | Predominantly oral-associated |

Permutation-based topological validation confirmed that the observed tissue-state organisation significantly exceeded random spatial structure (Table 7). All datasets demonstrated significantly elevated graph modularity and significantly reduced spatial entropy relative to permutation-null distributions (*p*_emp_=0.0099).

**Table 7:** Permutation-based topological validation. *R* = real non-permuted value, *P* = permutation-null mean, SD = standard deviation, CI = 95% confidence interval, and *p*_emp_ = empirical permutation *p*-value.

| Dataset | Metric | $R$ | $P \pm \text{SD}$ | CI | $p_{\text{emp}}$ |
| --- | --- | --- | --- | --- | --- |
| GSE241124 | Graph modularity | 0.0473 | $-0.0001 \pm 0.0009$ | $-0.0016-0.0017$ | 0.0099 |
| GSE241124 | Spatial entropy | 0.8770 | $0.8873 \pm 0.0004$ | $0.8864-0.8882$ | 0.0099 |
| GSE241124 | Mean local recurrence | 0.5738 | $0.5745 \pm 0.0005$ | $0.5736-0.5752$ | 0.9307 |
| GSE206621 | Graph modularity | 0.0582 | $0.0001 \pm 0.0007$ | $-0.0012-0.0015$ | 0.0099 |
| GSE206621 | Spatial entropy | 0.8552 | $0.8778 \pm 0.0003$ | $0.8772-0.8784$ | 0.0099 |
| GSE206621 | Mean local recurrence | 0.5596 | $0.5639 \pm 0.0004$ | $0.5630-0.5647$ | 1.0000 |
| GSE208253 | Graph modularity | 0.0416 | $0.0000 \pm 0.0008$ | $-0.0016-0.0017$ | 0.0099 |
| GSE208253 | Spatial entropy | 0.8696 | $0.8785 \pm 0.0003$ | $0.8778-0.8791$ | 0.0099 |
| GSE208253 | Mean local recurrence | 0.5673 | $0.5653 \pm 0.0004$ | $0.5645-0.5661$ | 0.0099 |
| GSE135805 | Graph modularity | 0.1061 | $-0.0003 \pm 0.0026$ | $-0.0056-0.0049$ | 0.0099 |
| GSE135805 | Spatial entropy | 0.6770 | $0.7883 \pm 0.0013$ | $0.7854-0.7912$ | 0.0099 |
| GSE135805 | Mean local recurrence | 0.6098 | $0.5815 \pm 0.0011$ | $0.5793-0.5837$ | 0.0099 |

In contrast, mean local recurrence exhibited mixed significance across datasets. GSE208253 and GSE135805 demonstrated significantly elevated recurrence relative to the permutation-null distributions, whereas GSE241124 (*p*_emp_=0.9307) and GSE206621 (*p*_emp_=1.0000) did not significantly differ from the null distributions despite preserving strong global topology. This distinction indicated that global graph organisation and local recurrence coherence capture complementary aspects of spatial tissue-state architecture.

Finally, summary statistics across datasets demonstrated consistently high state conservation entropy, strong domain coupling, moderate fragmentation, and preserved recurrence coherence across biological domains (Table 8).

**Table 8:** Summary statistics across datasets for spatial organisation metrics. *H*_*s*_ = mean state conservation entropy, *R*_*l*_ = mean local recurrence, and *H*_*p*_ = mean spatial entropy.

| Metric | Mean ± SD | Range (min–max) |
| --- | --- | --- |
| $H_s$ | $0.860 \pm 0.051$ | 0.763–0.900 |
| $C_{\text{global}}$ | $0.960 \pm 0.030$ | 0.940–0.994 |
| PFI | $0.422 \pm 0.022$ | 0.390–0.440 |
| $R_l$ | $0.641 \pm 0.027$ | 0.621–0.681 |
| $H_p$ | $1.726 \pm 0.083$ | 1.603–1.788 |

## 4 Discussion

By integrating heterogeneous spatial transcriptomic datasets using recurrence modelling, graph-based spatial learning, and tensor decomposition, the proposed framework identified conserved stromal, ECM, vascular, and inflammatory programmes shared across trauma, oral, and cardiovascular tissues. The findings support the hypothesis that biologically distinct tissue systems exhibit partially conserved spatial tissue-state organisation despite differences in tissue origin, pathology, and acquisition technology.

ECM remodelling and fibroblast-associated signalling emerged as the dominant conserved programmes across the latent tissue states. Tensor decomposition further identified ECM remodelling, fibroblast activation, and endothelial signalling as recurrent cross-domain pathway–state components linking trauma, oral, and cardiovascular tissues. These observations suggest that stromal and matrix-associated processes represent a conserved biological backbone underlying wound repair, chronic inflammation, tumour-associated remodelling, and cardiovascular tissue organisation.

The latent embedding analysis demonstrated substantial overlap between trauma and oral tissues within a shared latent manifold, consistent with their strong spatial coupling and similar tissue-state organisation. This behaviour is biologically plausible because oral inflammatory disease and wound repair both involve persistent tissue injury, ECM degradation, fibroblast activation, angiogenesis, and immune remodelling. Cardiovascular tissue demonstrated comparatively more compact latent organisation and lower pathological fragmentation, possibly reflecting the more structurally constrained architecture of cardiac tissue relative to the heterogeneous inflammatory microenvironments observed in trauma and oral disease. However, conserved stromal and vascular programmes remained evident across all domains.

Fuzzy tissue-state modelling revealed gradual transitions between neighbouring latent states rather than sharply separated clusters, supporting the presence of continuous biological progression and overlapping stromal–inflammatory microenvironments. Elevated fuzzy membership entropy across many spatial locations further suggests substantial biological heterogeneity and transitional tissue organisation during inflammation, fibrosis, and tissue remodelling.

The spatial tissue-state maps and recurrence analyses further demonstrated biologically meaningful differences in tissue organisation between disease contexts. Oral inflammatory tissue exhibited diffuse and fragmented spatial organisation, whereas oral squamous cell carcinoma demonstrated comparatively stronger regional coherence and recurrence reinforcement. These differences likely reflect heterogeneous inflammatory infiltration in chronic inflammatory lesions versus stronger compartmentalisation and stromal restructuring in tumour-associated microenvironments.

Permutation-based validation demonstrated significantly elevated graph modularity and reduced spatial entropy relative to null models, confirming that the latent tissue-state organisation was spatially structured and non-random. However, local recurrence significance varied across datasets, suggesting global graph topology and local recurrence density capture complementary aspects of tissue organisation at different spatial scales.

Methodologically, the proposed framework differs from conventional supervised disease-classification pipelines by focusing on unsupervised identification of conserved spatial tissue-state organisation across heterogeneous biological domains. The integration of recurrence modelling with graph-based spatial learning additionally enables continuous biological transitions and overlapping tissue states to be represented without imposing rigid discrete boundaries.

Several limitations should nevertheless be acknowledged. The cardiovascular dataset contributed fewer spatial observations than the trauma and oral datasets, potentially contributing to weaker cardiovascular representation within some latent tissue states. In addition, the current analysis focused primarily on transcriptomic spatial organisation and did not incorporate proteomic, imaging, metabolic, or longitudinal clinical information. The biological interpretation of latent tissue states also remains dependent on pathway annotation quality and embedding stability.

Despite these limitations, the study demonstrates the feasibility of integrating heterogeneous spatial transcriptomic datasets to identify conserved tissue-state organisation across clinically related biological systems. More broadly, the findings suggest that ECM remodelling, fibroblast activation, vascular adaptation, and inflammatory signalling represent recurrent spatial organisational principles linking trauma, oral, and cardiovascular tissues. Future work should extend the framework through multimodal spatial biology integration, single-cell transcriptomics, and larger clinically annotated cohorts to further evaluate the translational relevance of the conserved spatial programmes identified in this study.

## 5 Conclusion

The results demonstrated substantial latent overlap and strong spatial coupling between trauma and oral tissues, while cardiovascular tissue retained conserved but comparatively more spatially compact organisation. Spatial fragmentation, recurrence analysis, and permutation-based topological validation further confirmed that the observed tissue-state architecture was non-random and biologically structured.

These findings reveal conserved stromal and inflammatory programmes across trauma, oral pathology, and cardiovascular tissue, suggesting shared spatial regulatory mechanisms independent of tissue origin or disease aetiology.

This study demonstrates the utility of spatial AI frameworks for uncovering latent biological relationships across disparate pathological contexts, establishing a foundation for future multimodal spatial biology and translational research.

## Code Availability

The MATLAB package implementing the proposed framework is publicly available at the author‘s website under the heading “Spatial AI for cross-domain tissue-state analysis in trauma, oral, and cardiovascular pathology”: https://sites.google.com/view/tuan-d-pham/codes.

## Notes

https://www.ncbi.nlm.nih.gov/geo/query/acc.cgi?acc=GSE241124

https://www.ncbi.nlm.nih.gov/geo/query/acc.cgi?acc=GSE208253

https://www.ncbi.nlm.nih.gov/geo/query/acc.cgi?acc=GSE206621

https://www.ncbi.nlm.nih.gov/geo/query/acc.cgi?acc=GSE135805

### Competing Interest Statement

The authors have declared no competing interest.

## References

[1] Edmiston T, et al. Variation in global trauma care: a survey of 187 hospitals across 51 countries. BMJ Glob Health. 2025;10(11):e021784.

[2] Angjelova A, et al. Impact of periodontitis on endothelial risk dysfunction and oxidative stress improvement in patients with cardiovascular disease. J Clin Med. 2024;13(13):3781.

[3] Di X, Li Y, Wei J, Li T, Liao B. Targeting fibrosis: from molecular mechanisms to advanced therapies. Adv Sci. 2024:e2410416.

[4] Moretti L, Stalfort J, Barker TH, Abebayehu D. The interplay of fibroblasts, the extra-cellular matrix, and inflammation in scar formation. J Biol Chem. 2022;298(2):101530.

[5] Yoshimura A. Fibrosis: from mechanisms to novel treatments. Inflamm Regen. 2024;44(1):1.

[6] Li Q, Ouyang X, Lin J. The impact of periodontitis on vascular endothelial dysfunction. Front Cell Infect Microbiol. 2022;12:998313.

[7] Dhungana G, et al. Unveiling the molecular crosstalk between periodontal and cardiovascular diseases: a systematic review. Dent J. 2025;13(3):98.

[8] Foster DS, et al. Integrated spatial multiomics reveals fibroblast fate during tissue repair. Proc Natl Acad Sci USA. 2021;118(41):e2110025118.

[9] Younesi FS, et al. Fibroblast and myofibroblast activation in normal tissue repair and fibrosis. Nat Rev Mol Cell Biol. 2024;25:617–638.

[10] Williams CG, Lee HJ, Asatsuma T, Vento-Tormo R, Haque A. An introduction to spatial transcriptomics for biomedical research. Genome Med. 2022;14(1):68.

[11] Rao A, Barkley D, Franca GS, Yanai I. Exploring tissue architecture using spatial transcriptomics. Nature. 2021;596:211–220.

[12] Nguyen Q, et al. Spatial transcriptomics in human cardiac tissue. Int J Mol Sci. 2025;26(3):995.

[13] Cirillo MD, Mirdell R, Sjoberg F, Pham TD. Tensor decomposition for color image segmentation of burn wounds. Sci Rep. 2019;9:329.

[14] Wu Z, et al. Graph deep learning for the characterisation of tumour microenvironments from spatial protein profiles in tissue specimens. Nat Biomed Eng. 2022;6:1435– 1448.

[15] Long Y, et al. Spatially informed clustering, integration, and deconvolution of spatial transcriptomics with GraphST. Nat Commun. 2023;14(1):1155.

[16] Song T, Broadbent C, Kuang R. GNTD: reconstructing spatial transcriptomes with graph-guided neural tensor decomposition informed by spatial and functional relations. Nat Commun. 2023;14:8276.

[17] Kuppe C, et al. Spatial multi-omic map of human myocardial infarction. Nature. 2022;608(7924):766–777.

[18] Le DTP, Pham TD. Unveiling the role of artificial intelligence for wound assessment and wound healing prediction. Explor Med. 2023;4:589–611.

[19] Pham TD. Fuzzy recurrence exponents of subcellular-nanostructure dynamics in timelapse confocal imaging. IEEE Trans Nanobiosci. 2021;20(4):497–506.

[20] Pham TD. Recurrence eigenvalues of movements from brain signals. Brain Inform. 2021;8:22.

[21] Wang RH, Wang J, Li SC. Probabilistic tensor decomposition extracts better latent embeddings from single-cell multiomic data. Nucleic Acids Res. 2023;51(15):e81.

[22] Chen J, Han G, Xu A, Cai H. Identification of multidimensional regulatory modules through multi-graph matching with network constraints. IEEE Trans Biomed Eng. 2020;67(4):987–998.

[23] Yang X, Han G, Chen J, Cai H. Finding correlated patterns via high-order matching for multiple sourced biological data. IEEE Trans Biomed Eng. 2019;66(4):1017–1025.

[24] Pham TD, Yan H. Tensor decomposition of gait dynamics in Parkinson‘s disease. IEEE Trans Biomed Eng. 2018;65(8):1820–1827.

[25] Liu Z, et al. Spatial transcriptomics reveals that metabolic characteristics define the tumour immunosuppression microenvironment via iCAF transformation in oral squamous cell carcinoma. Int J Oral Sci. 2024;16:9.

[26] Benjamin K, et al. Multiscale topology classifies cells in subcellular spatial transcriptomics. Nature. 2024;630:943–949.

[27] Li D, et al. The lncRNA SNHG26 drives the inflammatory-to-proliferative state transition of keratinocyte progenitor cells during wound healing. Nat Commun. 2024;15:8637.

[28] Caetano AJ, et al. Spatially resolved transcriptomics reveals pro-inflammatory fibroblast involved in lymphocyte recruitment through CXCL8 and CXCL10. eLife. 2023;12:e81525.

[29] Arora R, et al. Spatial transcriptomics reveals distinct and conserved tumor core and edge architectures that predict survival and targeted therapy response. Nat Commun. 2023;14:5029.

[30] Egerstedt A, et al. Profiling of the plasma proteome across different stages of human heart failure. Nat Commun. 2019;10(1):5830.

[31] Yates AD, et al. Ensembl 2020. Nucleic Acids Res. 2020;48(D1):D682–D688.

[32] Tweedie S, et al. Genenames.org: the HGNC and VGNC resources in 2021. Nucleic Acids Res. 2021;49(D1):D939–D946.

[33] Subramanian A, et al. Gene set enrichment analysis: a knowledge-based approach for interpreting genome-wide expression profiles. Proc Natl Acad Sci USA. 2005;102(43):15545–15550.

[34] Bezdek JC, Ehrlich R, Full W. FCM: the fuzzy c-means clustering algorithm. Comput Geosci. 1984;10(2–3):191–203.

[35] Lin J. Divergence measures based on the Shannon entropy. IEEE Trans Inf Theory. 1991;37(1):145–151.

